# Skin colonization by circulating neoplastic clones in cutaneous T-cell lymphoma

**DOI:** 10.1101/703876

**Authors:** Aishwarya Iyer, Dylan Hennessey, Sandra O’Keefe, Jordan Patterson, Weiwei Wang, Gane Ka-Shu Wong, Robert Gniadecki

## Abstract

Mycosis fungoides (MF) is a mature T-cell lymphoma currently thought to develop primarily in the skin by a clonal expansion of a transformed, resident memory T-cell. However, this concept does not explain the key characteristics of MF such as the debut with multiple, widespread skin lesions or inability of skin directed therapies to provide cure. The testable inference of the mature T-cell theory is the clonality of MF with respect to all rearranged T-cell receptor (TCR) genes. Here we have used whole exome sequencing approach to detect and quantify TCRα, -β and -γ clonotypes in tumor cell clusters microdissected from MF lesions. This method allows us to calculate the tumor cell fraction of the sample and therefore an unequivocal identification of the TCR clonotypes as neoplastic. Analysis of TCR sequences from 29 patients with MF stage I-IV proved existence of multiple T-cell clones within the tumor cell fraction, with a considerable variation between patients and between lesions from the same patient (median 11 clones, range 2-80 clones/sample). We have also detected multiple neoplastic clones in the peripheral blood in all examined patients. Based on these findings we propose that circulating neoplastic T-cell clones continuously replenish the lesions of MF thus increasing their heterogeneity by a mechanism analogous to the consecutive tumor seeding. We hypothesize that circulating neoplastic clones might be a promising target for therapy and could be exploited as a potential biomarker in MF.

## Introduction

Cutaneous T-cell lymphomas (CTCL) are mature T-cell neoplasms, among which mycosis fungoides (MF) is the most common disease entity.^1^ MF presents initially as scaly, erythematous patches and plaques on skin, which may progress to tumors and disseminate to lymph nodes and other organs, such as the central nervous system.^2–4^ The pathogenesis of MF has been studied for decades as a model disease reflecting key characteristics of low-grade lymphomas such as progressive course and lack of curative treatments.

MF is believed to originate from the mature, memory, tissue-resident T-cells expressing skin homing markers CLA and CCR4.^5,6^ This straightforward hypothesis explains the affinity of MF to the skin and its low capacity to disseminate to extracutaneous sites. However, some clinical and molecular features of MF are incompatible with the model of the skin-resident memory T-cell as origin of MF. It is unexplainable why the disease usually starts multifocally in different areas of the skin rather than in a single site representing the location of the founding, transformed T-cell. Second, even profound depletion of lymphocytes in the skin (e.g. by electron beam radiation therapy or psoralen ultraviolet A therapy [PUVA]) almost never results in a cure but only in a short-term responses.^7–10^ Third, cells sharing molecular characteristics of malignant T-cells in MF have been found in the bone marrow of the patients years before the emergence of skin lesions of the disease^11^ and CTCL can be transmitted via the bone marrow transplant from asymptomatic donors^12,13^. Fourth, MF may share the common precursor with other lymphomas (e.g. Hodgkin lymphoma) that do not originate in the skin but primarily occupy extracutaneous sites, such as lymph nodes.^14^

These observations are more compatible with a scenario in which CTCL originate by hematogenous spread of precursor neoplastic cells to the skin niche.^15^ However, this concept was met with skepsis and resistance because analysis of the clonality of the T-cell receptor (TCR) seemed to strongly indicate that this disease is monoclonal and originates in the skin.^6,16^

TCR clonality assays have been the most powerful technique to dissect the pathogenesis of CTCL and other T-cell lymphomas.^17,18^ During T-cell development, the *V*, (*D*) and *J* gene segments of *TCRG*, *TCRB* and *TCRA* undergo sequential rearrangements producing unique CDR3 sequences which are retained in the mature T-cells. The diversity at CDR3 is further increased by insertions or deletions at *V(D)J* junctions and therefore those sequences constitute a unique signature of a given T-cell clone.^19^ Most research focused on *TCRG* because of its relatively small size and limited diversity and only recently on *TCRB* which is more diverse and has a unique property of allelic exclusion which simplifies data analysis.

We have recently shown that MF cells sampled from a plaque or a tumor may share the same TCRγ clonotype but exhibit different TCRβ and TCRα clonotypes ^20,21^. Since TCRγ loci (*TCRG*) rearrange before TCRβ (*TCRB)* and TCRα (*TCRA)* and the unique *TCRG* CDR3 sequences are inherited by the T-cells derived from those early clones, these findings are incompatible with the current model of mature T-cell as precursor of MF where all malignant cells would share an identical clonotype for all rearranged TCR genes (*TCRG*, *TCRB* and *TCRA)*.

Skin-resident T-cells do not recirculate but remain in the tissue where they can survive and proliferate without migration to the lymph nodes.^5,22,23^ Although atypical malignant T-cells are conspicuously absent in the blood in MF patients in the early stages and clonality assays are usually negative, it may be argued that standard TCRγ detection methods are rather insensitive for detecting rare clones in the background of highly diverse normal T-cells.^24^ Indeed, using more careful experimental approaches such as tumor fraction enrichment by laser capture microdissection and analysis of purified lymphocytes or mononuclear cells from the blood, some authors were able to find clonal, circulating cells even in early stages of disease development.^20,25–28^ Unfortunately, due to the difficulties in PCR amplification of *TCRB* and *TCRA* from genomic DNA these findings rely heavily on the analyses of TCRγ, which may give false positive results because of the low diversity of *TCRG*.

Here, we have applied the technique of TCR detection by whole exome sequencing ^21^ to revisit the hypothesis of circulating neoplastic cells in MF. By comparing TCR clonotypes in the skin and blood in patients with MF we reveal a complex pattern of recirculation of tumor subclones. We propose that substantial clonotypic heterogeneity of skin lesions in MF is caused by the mechanism of consecutive seeding of the skin niche by multiple subclones of neoplastic cells.

## Results

### *TCRB* sequencing of MF shows lesional and topological heterogeneity of malignant clonotypes

We have previously provided evidence that whole exome sequencing (WES) can be used for identifying neoplastic TCR clonotypes, i.e. the TCR CDR3 sequences specific to malignant T-cells in MF.^21^ Our method relies on the sequencing of CDR3 regions and quantification of the fraction of TCRα, -β and -γ clonotypes corresponding to the tumor cell percentage, thus filtering out the TCR sequences from the reactive, tumor-infiltrating T-cells. Using the same approach here, we have identified neoplastic clonotypes in 29 patients with MF using microdissected samples of 46 biopsies from cutaneous lesions (**Fig 1**, supplementary **Table S1**). To quantify clonogenic heterogeneity, we focused on *TCRB* which is sufficiently diverse to avoid the risk identical rearrangements of unrelated T-cell clones and which is rearranged on a single chromosome in >98% of all T-cells (allelic exclusion) thus unequivocally defining a T-cell clone.^29–32^ In a purely monoclonal disease, one can expect that the frequency of the single, dominant TCRβ clonotype matches the tumor cell fraction of the sample. However, in our 46 skin biopsies the tumor cell fraction significantly exceeded the frequency of the most abundant TCRβ clonotype which unequivocally proved clonotypic heterogeneity of MF. We identified a range of 2-80 TCRβ clonotypes per sample, which corresponded to the tumor cell fraction of the sample (i.e. neoplastic clonotypes) (**Fig 2A**, **Fig S1A**). On average, the most frequent (dominant) TCRβ clonotype comprised only 19.32% of the tumor fraction, which was similar to the values obtained in other studies using the PCR-based approach and NGS.^33,34^ The number of neoplastic TCRβ clonotypes correlated with the tumor cell fraction but not with the stage of the lesion (T1 plaque vs T2 tumor) which further supported the notion that those clonotypes represented true tumor clones and were not derived from infiltrating, reactive T-cells (**Fig 2B**).

**Figure 1:**
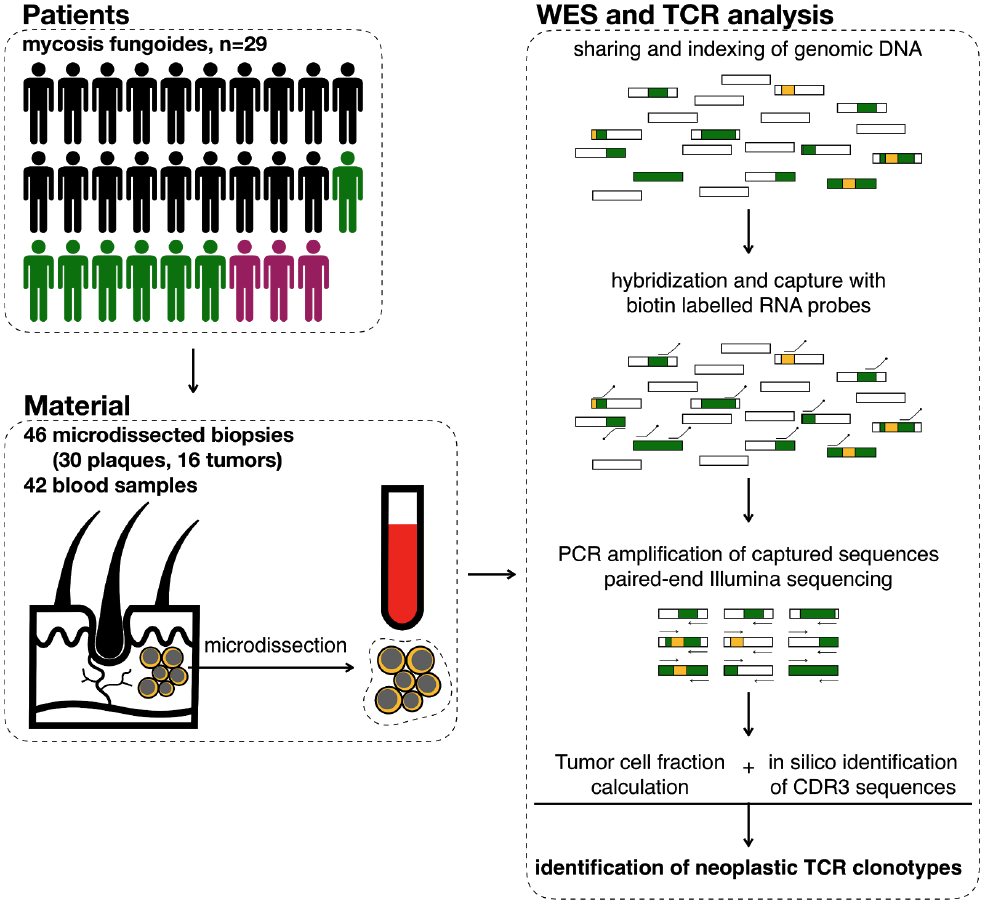
Schematic representation of sample collection and processing. 4mm single or multiple punch biopsies from and 10 ml of total blood was collected from 29 patients. In 7 patients we collected more than one biopsy (green silhouettes) whereas three patients were followed longitudinally with several biopsies and/or blood samples. The skin biopsies were cryosectioned and used for laser microdissection of clusters of tumour cells, which along with blood PBMC were processed for WES. Sequenced data was analyzed for identifying rearranged CDR3 sequences of *TCRA*, *TCRB* and *TCRG* and to determine tumor cell fraction. Rectangles represent DNA fragments, green areas are exons, yellow areas are rearranged TCR genes.

**Figure 2.**
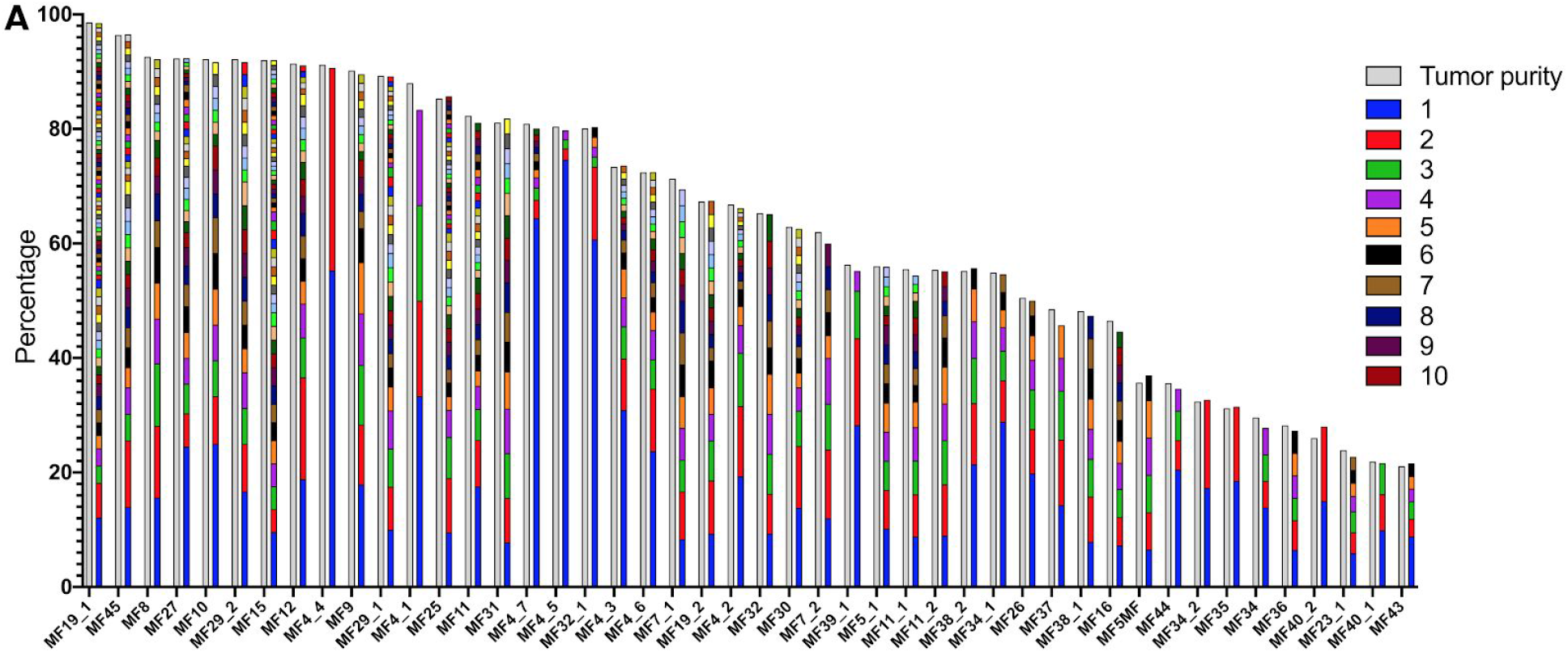
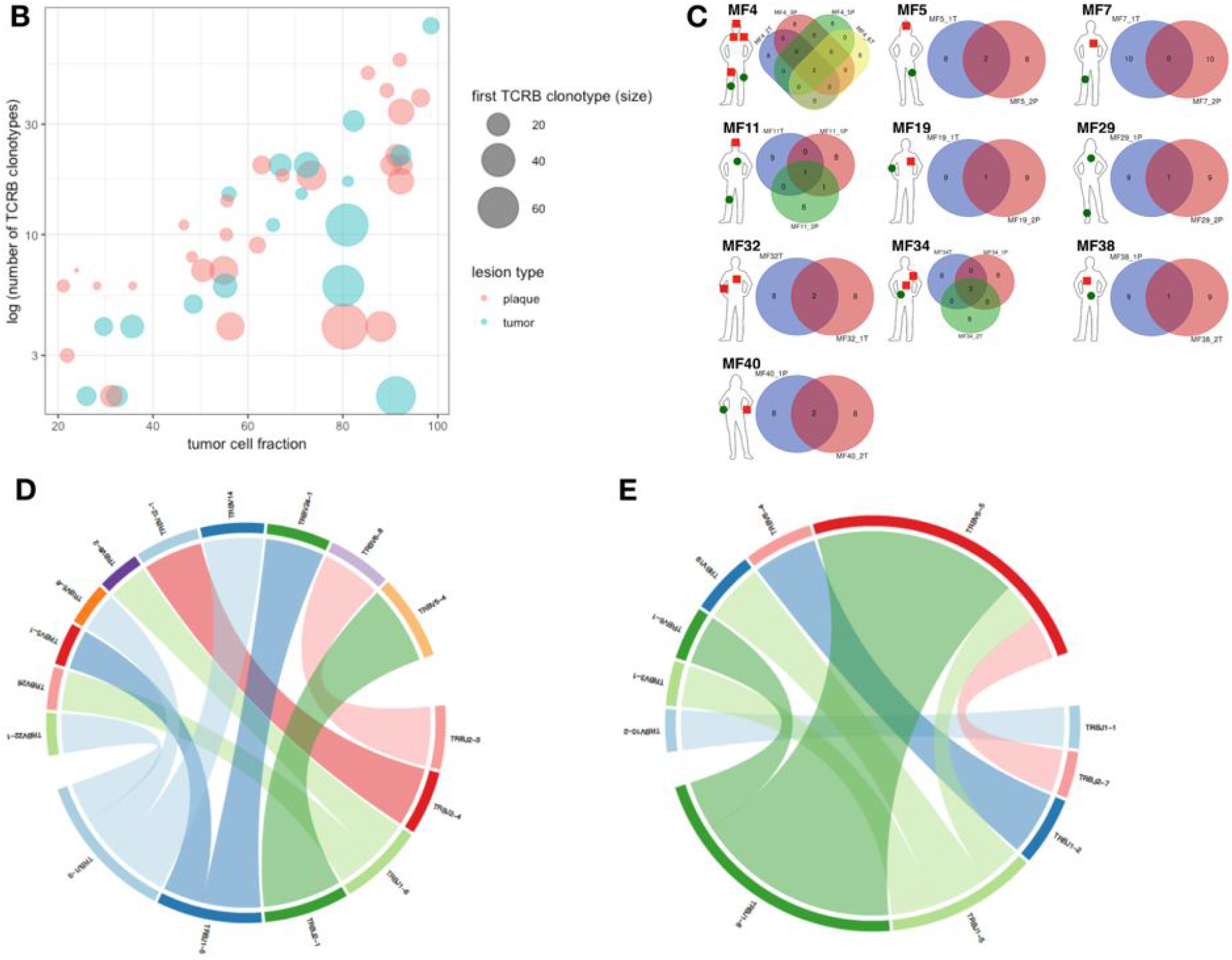
Clonotypic heterogeneity of skin lesions in MF. **A.** The tumor purity for the skin samples was estimated by the copy number aberration (CNA) data from the WES. The tumor cell fraction (grey bars) is plotted versus the cumulative frequency of the most abundant TCRβ clonotypes. The frequencies of the clonotypes are represented with stacked bars representing clonotypes from the most abundant (rank #1) to the least frequent. The ranks of clonotypes are color-coded as in the legend. **B.**Correlation between tumor cell fraction and the number of neoplastic clonotypes. Note that the clonotypic heterogeneity is not dependent on the stage of the lesion (tumor vs plaque). The size of the circle is proportional to the percentage of the most dominant (rank #1) TCRβ clonotype. **C.**Topological heterogeneity in MF. Venn diagrams illustrating the number of overlapping TCRβ clonotypes across different skin lesions. The location and type of the lesion is plotted for each patient (green circle - plaque, red square - tumor). **D-E**VJ combination diversity of TCRβ clonotypes in MF. The combinations of *VJ* genes of the neoplastic clonotypes is presented in D. MF32 and E. MF16.

In 10 patients we obtained biopsies from multiple skin lesions which enabled us to study clonotypic heterogeneity between different areas of the skin (**Fig 2C**, **Fig S5B-C**). In 8 patients we compared the biopsies from the plaque and the tumor (late lesion) (patients MF4, MF5, MF7, MF11, MF19, MF34, MF38 and MF40) and in 3 patients we compared lesions in the same stage (two plaques from MF29, stage T2 and two tumors from MF4 and MF32, stage T3). The heterogeneity of any single lesion was lower than the combined heterogeneity of all biopsies from the same patient combined, measured by the clonotype richness (number of different neoplastic TCRβ clonotypes) and Simpson index (the probability that two clonotypes, randomly drawn from the sample are different) (**supplementary Fig S1**). Surprisingly, the degree of overlap clonotypes between different lesions was very low (1-2 clonotypes) and in one case (MF7 tumor and plaque) we did not detect any shared clonotypes. Importantly, the shared clonotypes were not always the most frequent ones. Thus, extensive clonotypic heterogeneity was not detected only on the level of a single lesion, but also between different lesions (topological heterogeneity).

### Neoplastic clonotypes are frequently detected in the peripheral blood in MF

Having established that MF shows high clonotypic heterogeneity, both with regard to the composition of a single lesion and between different lesions, we realized that this heterogeneity could not be generated in the skin *in situ* because cells in the MF infiltrate do not express RAG1/2 and TdT, the key enzymes needed for TCR gene recombination (Ref^35^ and our unpublished data on RNAseq of MF). We considered the possibility that there is a pool of clonotypically heterogeneous neoplastic cells in the circulation which are able to seed the skin and undergo clonal expansion in the skin niche. Therefore, we investigated whether malignant T-cell clones could be found in the peripheral blood. For the consistency across samples, we assumed conservatively that the top 10 frequent TCR clonotypes from skin represent the true, tumor-related clonotypes. This assumption is based on the observation that the 10 most abundant TCRβ clonotypes contributed to up to 85% (95% upper confidence interval value) of the tumor-related clonotypes whereas the remaining clonotypes (ranked >10) had a low abundance (95% CI: 1.3%-1.5%,) and only contributed to 6%-15% (95% CI) of the total number of clonotypes. To provide a second layer of validation, we have also compared the *TCRA* and *TCRG* CDR3 sequences in the blood and the skin.

In 79% (15/19) of patients we have detected at least one shared TCRβ clonotype between the skin and the blood. The same number of patients had one or more common TCRɣ shared clonotype in the skin and blood, whereas 16 patients had circulating neoplastic TCRα clonotypes **(Fig 3A**, **Fig S2A and S3A**). Thus, all patients had at least one shared clonotype TCRα, -β or -γ between the skin and the blood at the time of sampling. The number of identical clonotypes in the skin and the blood was highest for TCRɑ (1-7 clonotypes, **Fig S2A**), probably due to the fact that a single clone of T-cells defined by a common TCRβ clonotype may comprise 1-3 different TCRα clonotypes (see supplementary **Fig S4**). Importantly, the neoplastic clonotypes were composed of multiple *V-J* gene combinations that indicated they originated at the stage of T-cell development (**Fig 2D, E**).

**Figure 3:**
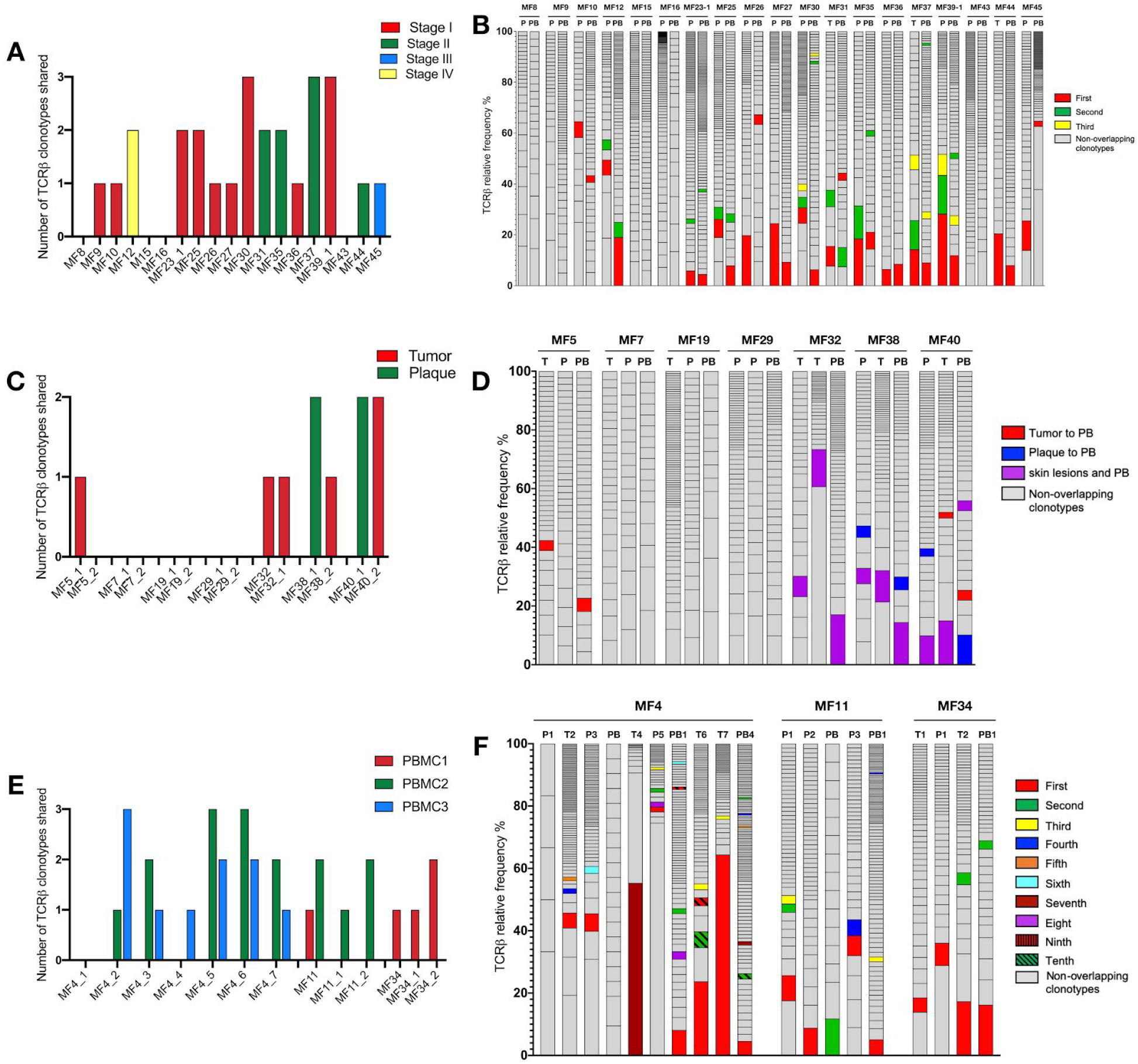
Detection of the neoplastic TCRβ clonotypes in the peripheral blood. The sequences of the TCRβ clonotypes in the blood that matched any of the top 10 neoplastic TCRβ clonotypes identified in the corresponding skin sample were identified to detect the neoplastic clonotypes in the circulation. The number and frequency of those shared neoplastic clonotypes are shown separately for the three groups of patients as defined in Fig 1: A, B: 19 patients with a single biopsy, C-F: patients with multiple skin biopsies, of whom in 7 patients the biopsies were obtained at a single time point (C, D) whereas 3 patients were sampled longitudinally (E, F). In B, D, F the first ranking shared clonotype in the skin is indicated in red and the subsequent shared clonotypes are color-coded as indicated in the legend. The non-overlapping clonotypes are indicated in gray.

We subsequently analysed whether the frequency of the clonotypes in the skin correlated with the frequency of those in the blood. The dominant (most abundant) TCRβ clonotype from the skin was identified in the blood in 8 patients and 6 of those clonotypes were also dominant in the blood (**Fig 3B**). The pattern was more complicated for TCRγ and TCRα, where dominant clonotypes could be detected in the blood and the skin in only 2 patients (MF36 and MF37) (**Fig S2B and S3B**).

Similar occurrence of tumor-derived clonotypes was detected in 10 patients in whom we analysed multiple skin biopsies. 7 patients were analyzed with paired biopsies (tumor and plaque) and for 3 patients multiple biopsies (between 3 and 7). All patients shared at least one clonotype (TCRβ, -α or -γ) between the blood and one or both skin biopsies (**Fig 3C**, **S2C, Fig S3C**) and the degree of clonotypic sharing was higher between the blood and the skin than between two skin biopsies (**Fig 3D**). No correlation was found between the number of shared clonotypes and the stage of the disease and the progression-free survival (Supplementary **Table 1**).

It has not escaped our attention that some clonotypes were shared between different samples (both skin and blood) from different patients, such as the CDR3 sequence CDNNNDMRF (TRAV16/TRAJ43) which was found among malignant clonotypes of patients 8 of 29 patients and CAASRGC_AKNIQYF (TRBV18/TRBJ2-4) that was found in 20 of 29 patients (see supplementary **Table S2**). We have previously noticed frequent clonotypic sharing between patients with MF and excluded laboratory error as a possible cause.^21^ We have also excluded the possibility that those sequences represented sequences parts of unrelated captured exomes, with the secondary verification using blastn and blastp that indicated the sequences to be TCR (data not shown).

### Temporal dynamics of malignant TCR clonotypes in the skin and blood

Clonotypic heterogeneity could be achieved by seeding the skin with malignant clones, a mechanism which is responsible for the formation of metastases in solid tumors.^36–39^. Metastatic seeding may occur by single cancer cells (in which case the metastasis represents a single subclone) or via continuous seeding when clusters of cancer cells transfer the entire heterogeneity of the primary tumor to the metastases^37–39^. The third seeding mechanism, referred to as the consecutive seeding, relies on sequential recruitment of neoplastic cells to the metastases and result in the metastatic lesions that only represent a fraction of the heterogeneity of the primary tumor (**supplementary Fig S6**).^39^ Clonotypic heterogeneity of skin lesions excluded the single-cell seeding events whereas the fact that we detected single neoplastic clonotypes in the blood rather than combinations of different clonotypes argued against continuous seeding. To further elucidate the mechanism of tumor seeding, we followed the malignant clonotypes in the skin and the blood in 3 patients (MF4, MF11, MF34) over a period of 9 to 22 months (**Fig 3E, F and Fig 4**). In each case we found neoplastic T-cell clonotypes in the blood, defined as those *TCRB* CDR3 sequences that were found in at least one skin biopsy. The number of circulating neoplastic clonotypes varied from 2 clonotypes in MF34 to 10 clonotypes in MF4. Circulating neoplastic clonotypes were not detected constantly in all blood samples and certain CDR3 sequences could be detected in the blood before occurrence in skin biopsies, e.g. GPGTRLLVLGERGLLGRGRGR_WVWFLRGVPGLCSGANVLTF or CASCPH_VSCRRP that were found in the blood of patient MF4 months before their detection in skin biopsies (**Fig 4A**). Together with the finding that a single skin lesion contained only a fraction of all possible neoplastic clonotypes, our data strongly supported the model of the development of the skin lesions in MF by consecutive seeding (**Fig 5**and **supplementary Fig S6**).

**Figure 4:**
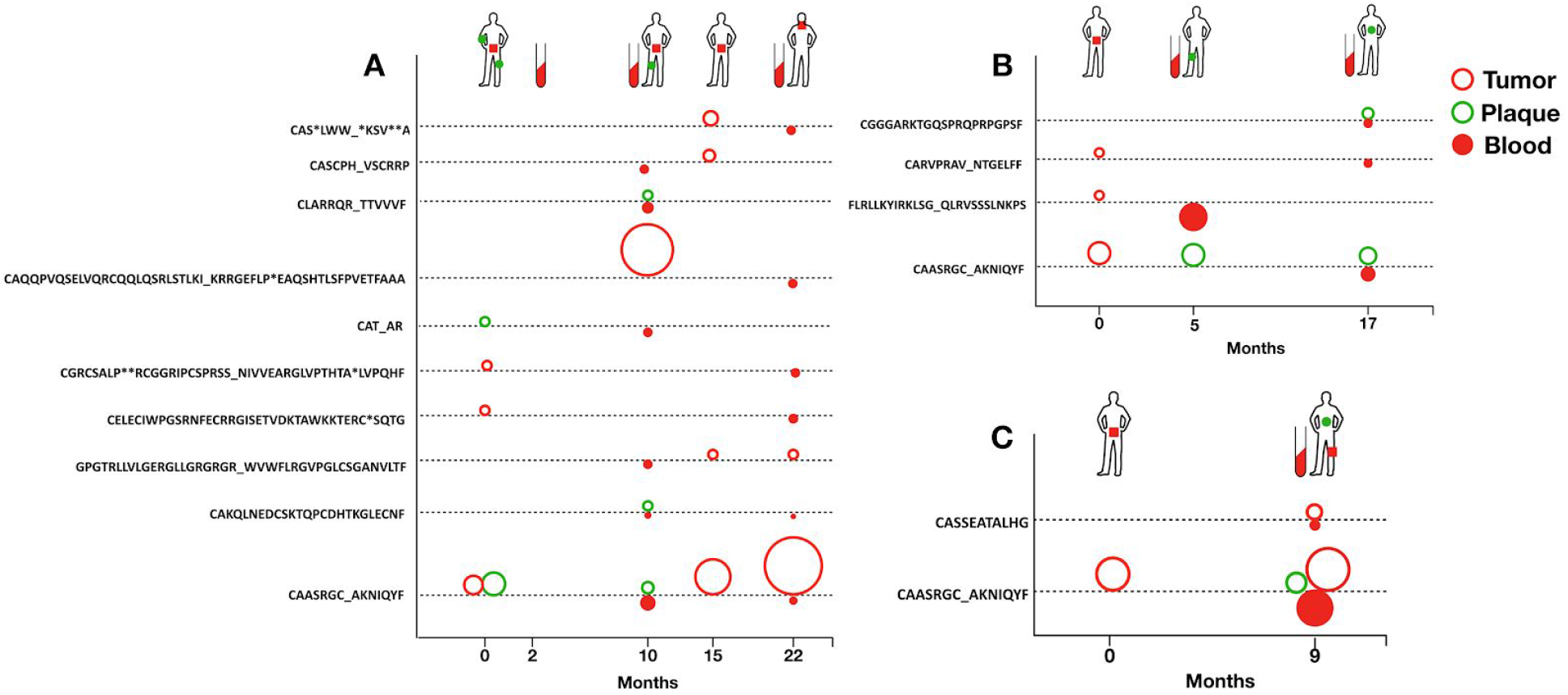
Dynamics of neoplastic clonotypes in the skin and the blood. Three patients were followed longitudinally with multiple skin biopsies and/or blood sampling. All shared neoplastic clonotypes were are plotted on the time axis for individual patients (A) MF4, (B) MF11 and (C) MF34. The location and type of the lesion is indicated for each patient on a silhouette (green circle - plaque, red square - tumor). Each dotted line corresponds to a single, shared clonotype of the indicated amino acid sequence of CDR3β. Circles above the line are skin clonotypes (open red-tumor, open green-plaque) whereas the solid red circles below the line are the clonotypes detected in the blood. The size of the dot is proportional to the frequency of the shared clonotype in the sample.

**Figure 5.**
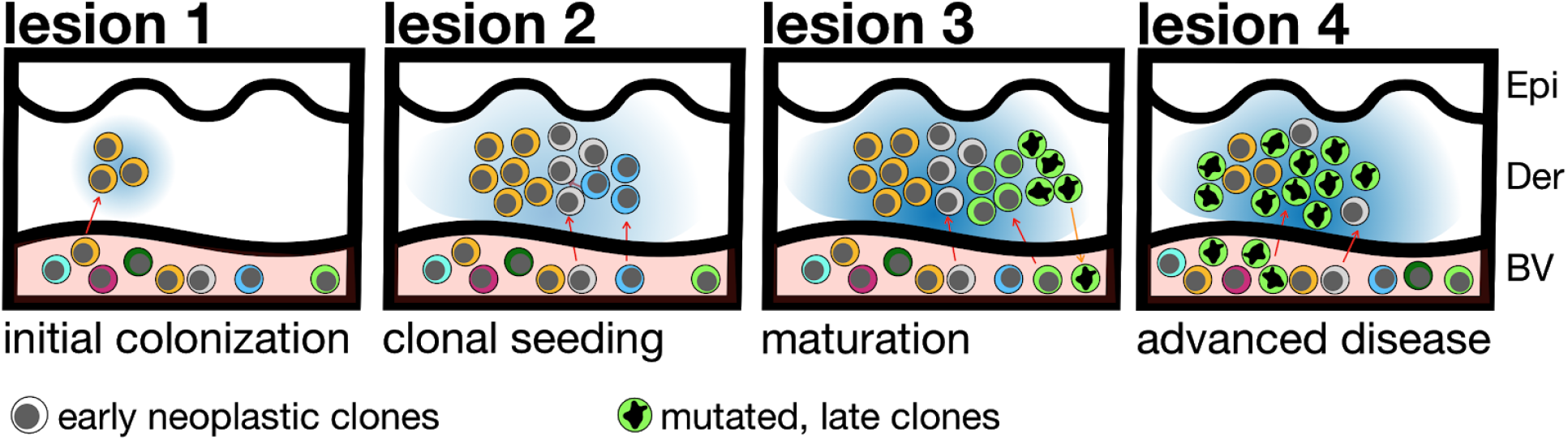
The hypothesis of tumor seeding in the pathogenesis of mycosis fungoides. Even in the early stages of the disease the patients have circulating neoplastic T-cell clones in peripheral blood. Early lesions are initiated by the pioneer clones and create a niche (blue-shaded area) that facilitates seeding of this area of the skin (“lesion 1”) with subsequent clones (“lesion 2”) (consecutive seeding model^39^ and Fig S6). The clonal composition of different lesions may differ (“lesion 3”) due to the stochastic nature of cancer seeding. Some clones may have a higher proliferation capacity in the skin and may overgrow other clones (green clone in “lesion 3”, further mutate, re-enter the circulation (orange arrow) and re-seed another area of the skin (lesion 4). The figures represent symbolically the structure of the skin with the epidermis (Epi), dermis (Der) and a pink-shaded blood vessel (BV). Different clones of neoplastic T-cells are marked with different colors.

## Discussion

The major finding of this study was that lesions of mycosis fungoides comprise a highly diverse collection of malignant T-cell clones which percolate between the skin and the blood. We propose that consecutive seeding of circulating neoplastic T-cell subclones in the skin is a likely mechanism of growth and evolution of the lesions in this cutaneous lymphoma.

Difficulties in distinguishing between the CDR3 sequences in malignant T-cells from those of normal, reactive T-cells has been the major limitation of TCR analyses. Several indirect approaches have been used to mitigate this problem, such as setting arbitrary thresholds for clonotypic frequencies or surrogate measures of tumor cell fraction as a ratio of the frequency of the most abundant TCRβ clonotype to the sum of all TCRβ clonotypes.^33^ Obviously, both approaches assume that all tumor cells are clonal (i.e. share a single TCRβ clonotype). This assumption is a cornerstone of the theory of CTCL as a neoplasm of the mature T-cell but has not rigorously been tested experimentally. Experimental findings suggesting clonotypic heterogeneity in CTCL^40^ have either been ignored or attributed to contamination by clonal inflammatory T-cells^41^, activation of skin resident T-cells by superantigens^40^ or age-dependent clonal expansion^42–45^. We were able to solve this methodological problem by calculating the tumor cell fraction in the sample by WES and thus directly enumerate the clonotypes derived from neoplastic T-cells^21^. Our method relies on DNA sequencing which eliminates a potential error due to aberrant TCR expression in cancer cells. With this approach we confirmed our previous findings^20^ demonstrating clonal heterogeneity in MF.

We have found a large variation in the clonotypic richness (number of different TCRβ clonotypes) in MF lesions, ranging from 2 up to 80 distinct clonotypes. In most samples, the frequency of the most abundant TCRβ clonotype has been relatively high (19.4%) which is well above the usual 15% threshold accepted by many authors as a hallmark of monoclonality. This is reflected by a relatively low Simpson index (a probability that a random draw from the pool of clonotypes yields two different clonotypes; median 7% 1st and 3rd quartile 3.2% - 10.7%) (**Fig S1**). Thus, even consecutive draws from the pool of clonotypes of a given lesion are most likely to yield identical clonotypes, which can be misinterpreted as monoclonality (e.g. 10 consecutive draws yields identical results in ∼50% of cases). In 33% of biopsies, the frequency of the first TCRβ clone was lower than 10%, which corresponds well to the observed proportion of cases of MF in which monoclonality cannot be detected by standard assays.^46^

Clonotypic heterogeneity could not be attributed to secondary somatic mutations within the already assembled CDR3 region because we detected that the rearranged *TCRB* were composed by numerous combinations of various VJ segments. Another argument against extensive secondary mutations within TCRB comes from the quantification of the ratio between TCRα and TCRβ clonotypes. In a population of normal memory T-cells the *TCRA/TCRB* is between 2 and 3 because *TCRA* rearrangements are usually biallelic and there are occasional secondary rearrangements of this locus.^47^ Mutations in *TCRB* would result in an increase in the number of unique TCRβ sequences and thus in a decrease in the *TCRA*/*TCRB* ratio, which was not the case in our material (*TCRA/TCRB* was 2.76). Third, we have detected a higher than expected overlap between clonotypes between patients (Supplementary **Table S2**) which argues against the influence of random mutations.

Conceptually, the clonotypic heterogeneity described here is different from the mutational subclonal heterogeneity, because it cannot be generated continuously in the tumor but only in the time span when RAG1/2 recombinases are active (i.e. at the level of immature T-cell precursor). It has been suggested though, that multiple malignant clones can be generated from the pool of normal, reactive lymphocytes in the skin undergoing malignant transformation.^40^ However, such a mechanism would have to operate at an unprecedented efficiency to generate hundreds of cancer clones simultaneously in different areas of the skin and is therefore unlikely. The most likely mechanism is by accrual of malignant T-cells clones from the circulation to the skin. Several lines of indirect evidence support such a hypothesis. First, we were able to detect malignant TCR clonotypes in the blood. It has long been known that TCRγ clonotypes identical to those in lesional skin could be detected in the blood in MF in 5%-10% of patients in early stage disease without any prognostic impact.^48–51^ We have found that the presence of circulating malignant clones is a rule rather than an exception; at least one malignant TCRβ or TCRα clonotype was present in the peripheral blood in all examined patients. Although only a small fraction of the entire pool of neoplastic clonotypes could be detected in the blood, the chances of detecting TCRβ clonotypes increased by repeated sampling, probably because of their very low frequency compared to the background of normal T-cells. Second, the neoplastic clonotypes were found in the blood even in the very early stages of the disease, they were not correlated with the stage of the disease nor they always represented the dominant clonotype in the skin. Therefore, those circulating clones could not solely represent subclinical leukemization due to disease progression. Third, clonotypic overlap (the number of shared TCRɑ, -β or -ɣ clonotypes) was higher between the skin and the peripheral blood than between discrete skin lesions.

Taken together, these observations are compatible with the presence of a pool of very diverse neoplastic T-cell clones in the peripheral blood that may seed at different frequency to the skin where they develop further into the lesions of MF (**Fig 5** **and S6**). Our findings are compatible with the model of consecutive seeding in which only a fraction of all neoplastic clones transfers the diversity to the developing lesions of lymphoma. Previous modeling of consecutive metastatic seeding events revealed that each lesion is likely to be funded by 10-150 cancer clones^39^, a number which is in the same range as the 6-20 clonotypes (1st-3rd quartiles) detected by us in MF. Although not investigated here directly, we hypothesize that neoplastic clones do not migrate unidirectionally from the blood to the skin. Recent findings that downregulation of CD69 enables skin resident memory cells to exit the tissue and to recirculate^52^ indicates that neoplastic cells in MF should also be able to re-enter the circulation and contribute to the pool of circulating neoplastic clones. This mechanism is reminiscent of the phenomenon of tumor self-seeding^36,53^ where circulating metastatic cells colonize the primary tumor. Tumor self-seeding is enhanced by changes in the tissue niche occupied by cancer, which makes the niche more receptive for the subsequent entry of the waves of circulating tumor cells.^53^ In MF, the source of those dermotropic neoplastic T-cell clones remains unknown, although there is some evidence that bone marrow may play such a role.^11–13^ Research showed that the bone marrow niche provides shelter for different types of malignant cells (e.g. breast or prostate cancers) and that the bone marrow pool of cancer might be responsible for relapses after therapy.^54,55^

The proposed mechanism of tumor self-seeding in the pathogenesis of MF may have several practical implications. Circulating neoplastic T-cell clones may constitute an interesting target for therapy. It is tempting to speculate that mogamulizumab^56^ which blocks CCR4, an essential cutaneous homing receptor^57^, exerts it therapeutic efficacy via inhibition of skin seeding by circulating cancer clones. Identification of TCRβ clonotypes could also be used diagnostically in CTCL. Interestingly, we have found significant interindividual overlap in TCRβ clonotypic sequences, dramatically exceeding the frequency of shared, public clonotypes in healthy individual. Although the mechanism of clonotypic sharing remains elusive, the common *TCRB* CDR3 sequences could easily, quantitatively and cost-effectively be detected with PCR-based techniques and used for the diagnosis and monitoring of therapy in CTCL.

## Materials and Methods

### Sample collection and storage

Ethical approval was obtained from the Health Research Ethics Board of Alberta, Cancer Committee HREBA.CC-16-0820-REN1. After informed consent, 4mm punch skin biopsies were collected from patients and embedded in optimal cutting temperature (OCT) medium stored at −80°C. 10 ml of blood was collected into EDTA tube, peripheral blood mononuclear cells (PBMC) were isolated by Ficoll centrifugation which were resuspended in 50% of Dulbecco’s modified eagle medium (DMEM) (cat# 11965-084) (Thermo Fisher Scientific, Massachusetts, United States), 40% Fetal bovine serum (FBS) (cat# 16000044) (Thermo Fisher Scientific) and 10% Dimethyl Sulfoxide (DMSO) (cat# 20688) (Thermo Fisher Scientific) and frozen in liquid nitrogen until further use. Before DNA isolation the PBMC cells were thawed, resuspended in Roswell Park Memorial Institute (RPMI) 1640 medium with 10% FBS. DNA and RNA was isolated with Trizol reagent (cat# 15596026) (Invitrogen, Carlsbad, California, United States).

### Cryosectioning, laser capture microdissection (LCM) and sample preparation for whole exome sequencing (WES)

Skin biopsies were cryosectioned and prepared for whole exome sequencing according to the previously reported protocol^21^. NEBNext^®^ Ultra^TM^ II DNA library prep kit for illumina (cat# E7645S) (New England Biolabs, Massachusetts, United States) was used for preparing the samples for sequencing and SSELXT Human All exon V6 +UTR probes (Agilent Technologies, California, United State) were used for the exome capture. The DNA libraries were sequenced on an Illumina HiSeq 1500 sequencer using paired-end (PE) 150 kit (cat# PE-402-4002) (Hiseq PE rapid cluster kit V2) or NovaSeq 6000 S4 reagent kit 300 cycles (cat# 20012866).

### Data analysis

The fastq files were analyzed using MiXCR (version 2.10.0) to identify the TCR clonotypes. For WES data, partial reads were filtered out as these might be the captures of only V or J sequences. The reads were processed using the GATK4 (version 4.0.10) generic data-preprocessing workflow, then analyzed with Titan (version 1.20.1) to determine copy number aberration and tumor cell fraction (TCF) using the hg38 Human reference genome. The tcR package for R was used to calculate the overlapping clones.

The number of neoplastic TCRβ clonotypes (*n*_*β*_) is calculated to satisfy the following formula: 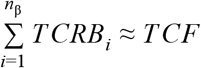, where *TCRB*_*i*_ is the percentage of the TCRβ clonotype of *i*-rank (the rank *i*=1 being the most abundant, dominant clonotype) and TCF is the tumor cell fraction in the sample calculated from WES. We assumed that the proportion of malignant T-cells cells with >1 rearranged TCRB is negligible (allelic exclusion) and therefore the number of neoplastic TCRβ clonotypes n_β_ is equal to the number of malignant T-cell clones. Although we have also computed the number of neoplastic clonotypes for TCRα (n_α_) and TCRγ (n_γ_), those values cannot directly be used for estimating the number of malignant clones because *TCRG* and *TCRA* often, but not always, rearrange on both chromosomes and *TCRA* may re-rearrange producing >2 clonotypes for each TCR β clonotype.

## Data availability

The data is currently being submitted to dbGAP. For the peer review process the data can be accessed via the corresponding author.

## Acknowledgement

We would like to acknowledge Dr. Thomas Salopek, Mrs. Rachel Doucet and the nursing staff of Edmonton Kaye clinic for their help in sample collection. Dr. Hanne Fogh provided excellent care to the patients from the Copenhagen center and helped to collect clinical data and Mrs Vibeke Pless contribution to the collection of samples. This study was supported by grants from the following sources: Canadian Dermatology Foundation (CDF RES0035718), University of Alberta, University Hospital Foundation (University of Alberta), Bispebjerg Hospital (Copenhagen, Denmark) unrestricted research grant to R.G., and Danish Cancer Society (Kræftens Bekæmpelse R124-A7592 Rp12350).

## Authors contribution

AI designed, analyzed the data and wrote the manuscript. AI and SO performed the experiments. JP and DH performed CNA analysis and tumor cell fraction calculations. WW and GW provided input with the technical aspects of the experiments and edited the manuscript. RG supervised the experiments, data analysis and edited the manuscript. All authors approved the final version of this paper.

## References

1. Swerdlow SH, Campo E, Pileri SA, et al. The 2016 revision of the World Health Organization classification of lymphoid neoplasms. Blood. 2016;127(20):2375–2390.

2. Olsen E, Vonderheid E, Pimpinelli N, et al. Revisions to the staging and classification of mycosis fungoides and Sezary syndrome: a proposal of the International Society for Cutaneous Lymphomas (ISCL) and the cutaneous lymphoma task force of the European Organization of Research and Treatment of Cancer (EORTC). Blood. 2007;110(6):1713–1722.

3. Kim YH, Willemze R, Pimpinelli N, et al. TNM classification system for primary cutaneous lymphomas other than mycosis fungoides and Sezary syndrome: a proposal of the International Society for Cutaneous Lymphomas (ISCL) and the Cutaneous Lymphoma Task Force of the European Organization of Research and Treatment of Cancer (EORTC). Blood. 2007;110(2):479–484.

4. Willemze R. WHO-EORTC classification for cutaneous lymphomas. Blood. 2005;105(10):3768–3785.

5. Clark RA, Watanabe R, Teague JE, et al. Skin effector memory T cells do not recirculate and provide immune protection in alemtuzumab-treated CTCL patients. Sci. Transl. Med. 2012;4(117):117ra7.

6. Campbell JJ, Clark RA, Watanabe R, Kupper TS. Sezary syndrome and mycosis fungoides arise from distinct T-cell subsets: a biologic rationale for their distinct clinical behaviors. Blood. 2010;116(5):767–771.

7. Whittaker S, Ortiz P, Dummer R, et al. Efficacy and safety of bexarotene combined with psoralen-ultraviolet A (PUVA) compared with PUVA treatment alone in stage IB-IIA mycosis fungoides: final results from the EORTC Cutaneous Lymphoma Task Force phase III randomized clinical trial 21011 (NCT00. British Journal of Dermatology. 2012;167(3):678–687.

8. Kamstrup MR, Gniadecki R, Iversen L, et al. Low-dose (10-Gy) total skin electron beam therapy for cutaneous T-cell lymphoma: an open clinical study and pooled data analysis. Int. J. Radiat. Oncol. Biol. Phys. 2015;92(1):138–143.

9. Chowdhary M, Chhabra AM, Kharod S, Marwaha G. Total Skin Electron Beam Therapy in the Treatment of Mycosis Fungoides: A Review of Conventional and Low-Dose Regimens. Clin. Lymphoma Myeloma Leuk. 2016;16(12):662–671.

10. Vieyra-Garcia P, Fink-Puches R, Porkert S, et al. Evaluation of Low-Dose, Low-Frequency Oral Psoralen-UV-A Treatment With or Without Maintenance on Early-Stage Mycosis Fungoides: A Randomized Clinical Trial. JAMA Dermatol. 2019;

11. Gniadecki R, Lukowsky A, Rossen K, et al. Bone marrow precursor of extranodal T-cell lymphoma. Blood. 2003;102(10):3797–3799.

12. Berg KD, Brinster NK, Huhn KM, et al. Transmission of a T-cell lymphoma by allogeneic bone marrow transplantation. N. Engl. J. Med. 2001;345(20):1458–1463.

13. Fahy CMR, Fortune A, Quinn F, et al. Development of mycosis fungoides after bone marrow transplantation for chronic myeloid leukaemia: transmission from an allogeneic donor. Br. J. Dermatol. 2014;170(2):462–467.

14. Davis TH, Morton CC, Miller-Cassman R, Balk SP, Kadin ME. Hodgkin’s disease, lymphomatoid papulosis, and cutaneous T-cell lymphoma derived from a common T-cell clone. N. Engl. J. Med. 1992;326(17):1115–1122.

15. Kamstrup MR, Gniadecki R, Skovgaard GL. Putative cancer stem cells in cutaneous malignancies. Exp. Dermatol. 2007;16(4):297–301.

16. Delfau-Larue M, Petrella T, Lahet C, et al. Value of clonality studies of cutaneous T lymphocytes in the diagnosis and follow-up of patients with mycosis fungoides. The Journal of Pathology. 1998;184(2):185–190.

17. Jones D, Duvic M. The current state and future of clonality studies in mycosis fungoides. J. Invest. Dermatol. 2003;121(3):ix–x.

18. Mahe E, Pugh T, Kamel-Reid S. T cell clonality assessment: past, present and future. J. Clin. Pathol. 2018;71(3):195–200.

19. Krangel MS. Mechanics of T cell receptor gene rearrangement. Curr. Opin. Immunol. 2009;21(2):133–139.

20. Hamrouni A, Fogh H, Zak ZL, Odum N, Gniadecki R. Clonotypic diversity of the T-cell receptor corroborates the immature precursor origin of cutaneous T-cell lymphoma. Clin. Cancer Res. 2019;

21. Iyer A, Hennessey D, O’Keefe S, et al. Clonotypic heterogeneity in cutaneous T-cell lymphoma (mycosis fungoides) revealed by comprehensive whole-exome sequencing. Blood Adv. 2019;3(7):1175–1184.

22. Boyman O, Hefti HP, Conrad C, et al. Spontaneous Development of Psoriasis in a New Animal Model Shows an Essential Role for Resident T Cells and Tumor Necrosis Factor-α. The Journal of Experimental Medicine. 2004;199(5):731–736.

23. Watanabe R, Gehad A, Yang C, et al. Human skin is protected by four functionally and phenotypically discrete populations of resident and recirculating memory T cells. Sci. Transl. Med. 2015;7(279):279ra39.

24. Langerak AW, Molina TJ, Lavender FL, et al. Polymerase chain reaction-based clonality testing in tissue samples with reactive lymphoproliferations: usefulness and pitfalls. A report of the BIOMED-2 Concerted Action BMH4-CT98-3936. Leukemia. 2007;21(2):222–229.

25. Marcus Muche J, Karenko L, Gellrich S, et al. Cellular coincidence of clonal T cell receptor rearrangements and complex clonal chromosomal aberrations-a hallmark of malignancy in cutaneous T cell lymphoma. J. Invest. Dermatol. 2004;122(3):574–578.

26. Muche JM, Lukowsky A, Heim J, et al. Demonstration of frequent occurrence of clonal T cells in the peripheral blood but not in the skin of patients with small plaque parapsoriasis. Blood. 1999;94(4):1409–1417.

27. Beylot-Barry M, Sibaud V, Thiebaut R, et al. Evidence that an identical T cell clone in skin and peripheral blood lymphocytes is an independent prognostic factor in primary cutaneous T cell lymphomas. J. Invest. Dermatol. 2001;117(4):920–926.

28. Fraser-Andrews EA, Woolford AJ, Russell-Jones R, Whittaker SJ, Seed PT. Detection of a Peripheral Blood T Cell Clone is an Independent Prognostic Marker in Mycosis Fungoides. Journal of Investigative Dermatology. 2000;114(1):117–121.

29. Pogorelyy MV, Elhanati Y, Marcou Q, et al. Persisting fetal clonotypes influence the structure and overlap of adult human T cell receptor repertoires. PLoS Comput. Biol. 2017;13(7):e1005572.

30. Dupic T, Marcou Q, Walczak AM, Mora T. Genesis of the αβ T-cell receptor. PLoS Comput. Biol. 2019;15(3):e1006874.

31. Khor B, Sleckman BP. Allelic exclusion at the TCRbeta locus. Curr. Opin. Immunol. 2002;14(2):230–234.

32. Mora T, Walczak AM. Quantifying lymphocyte receptor diversity.

33. de Masson A, O’Malley JT, Elco CP, et al. High-throughput sequencing of the T cell receptor β gene identifies aggressive early-stage mycosis fungoides. Sci. Transl. Med. 2018;10(440.)

34. Kirsch IL, Sherwood A, Williamson D, et al. The utility of HTS TCR analyses in the management of patients with T-cell malignancies. Journal of Clinical Oncology. 2014;32(15_suppl):8565–8565.

35. Döbbeling U, Dummer R, Hess Schmid M, Burg G. Lack of expression of the recombination activating genes RAG-1 and RAG-2 in cutaneous T-cell lymphoma: pathogenic implications. Clin. Exp. Dermatol. 1997;22(5):230–233.

36. Kim M-Y, Oskarsson T, Acharyya S, et al. Tumor Self-Seeding by Circulating Cancer Cells. Cell. 2009;139(7):1315–1326.

37. Aceto N, Bardia A, Miyamoto DT, et al. Circulating tumor cell clusters are oligoclonal precursors of breast cancer metastasis. Cell. 2014;158(5):1110–1122.

38. Cleary AS, Leonard TL, Gestl SA, Gunther EJ. Tumour cell heterogeneity maintained by cooperating subclones in Wnt-driven mammary cancers. Nature. 2014;508(7494):113–117.

39. Heyde A, Reiter JG, Naxerova K, Nowak MA. Consecutive seeding and transfer of genetic diversity in metastasis. Proceedings of the National Academy of Sciences. 2019;116(28):14129–14137.

40. Vega F, Luthra R, Medeiros LJ, et al. Clonal heterogeneity in mycosis fungoides and its relationship to clinical course. Blood. 2002;100(9):3369–3373.

41. Kolowos W, Herrmann M, Ponner BB, et al. Detection of restricted junctional diversity of peripheral T cells in SLE patients by spectratyping. Lupus. 1997;6(9):701–707.

42. Gniadecki R, Lukowsky A. Monoclonal T-cell dyscrasia of undetermined significance associated with recalcitrant erythroderma. Arch. Dermatol. 2005;141(3):361–367.

43. Wack A, Cossarizza A, Heltai S, et al. Age-related modifications of the human alphabeta T cell repertoire due to different clonal expansions in the CD4+ and CD8+ subsets. Int. Immunol. 1998;10(9):1281–1288.

44. Posnett DN, Sinha R, Kabak S, Russo C. Clonal populations of T cells in normal elderly humans: the T cell equivalent to “benign monoclonal gammapathy.” J. Exp. Med. 1994;179(2):609–618.

45. Muche JM, Sterry W, Gellrich S, et al. Peripheral blood T-cell clonality in mycosis fungoides and nonlymphoma controls. Diagn. Mol. Pathol. 2003;12(3):142–150.

46. Dereure O, Levi E, Vonderheid EC, Kadin ME. Improved sensitivity of T-cell clonality detection in mycosis fungoides by hand microdissection and heteroduplex analysis. Arch. Dermatol. 2003;139(12):1571–1575.

47. Hamrouni A, Aublin A, Guillaume P, Maryanski JL. T cell receptor gene rearrangement lineage analysis reveals clues for the origin of highly restricted antigen-specific repertoires. J. Exp. Med. 2003;197(5):601–614.

48. Muche JM, Marcus Muche J, Lukowsky A, et al. Peripheral Blood T Cell Clonality in Mycosis Fungoides –An Independent Prognostic Marker? Journal of Investigative Dermatology. 2000;115(3):504–505.

49. Muche JM, Marcus Muche J, Sterry W, et al. Peripheral Blood T-Cell Clonality in Mycosis Fungoides and Nonlymphoma Controls. Diagnostic Molecular Pathology. 2003;12(3):142–150.

50. Scarisbrick JJ, Hodak E, Bagot M, et al. Developments in the understanding of blood involvement and stage in mycosis fungoides/Sezary syndrome. Eur. J. Cancer. 2018;101:278–280.

51. Talpur R, Singh L, Daulat S, et al. Long-term outcomes of 1,263 patients with mycosis fungoides and Sézary syndrome from 1982 to 2009. Clin. Cancer Res. 2012;18(18):5051–5060.

52. Klicznik MM, Morawski PA, Höllbacher B, et al. Human CD4CD103 cutaneous resident memory T cells are found in the circulation of healthy individuals. Sci Immunol. 2019;4(37.):

53. Massagué J, Obenauf AC. Metastatic colonization by circulating tumour cells. Nature. 2016;529(7586):298–306.

54. Shiozawa Y, Pedersen EA, Havens AM, et al. Human prostate cancer metastases target the hematopoietic stem cell niche to establish footholds in mouse bone marrow. J. Clin. Invest. 2011;121(4):1298–1312.

55. Zhang XH-F, Jin X, Malladi S, et al. Selection of bone metastasis seeds by mesenchymal signals in the primary tumor stroma. Cell. 2013;154(5):1060–1073.

56. Kim YH, Bagot M, Pinter-Brown L, et al. Mogamulizumab versus vorinostat in previously treated cutaneous T-cell lymphoma (MAVORIC): an international, open-label, randomised, controlled phase 3 trial. Lancet Oncol. 2018;19(9):1192–1204.

57. Soler D, Humphreys TL, Spinola SM, Campbell JJ. CCR4 versus CCR10 in human cutaneous TH lymphocyte trafficking. Blood. 2003;101(5):1677–1682.

