## supplementary file for "Skin colonization by circulating neoplastic clones in cutaneous T-cell lymphoma"

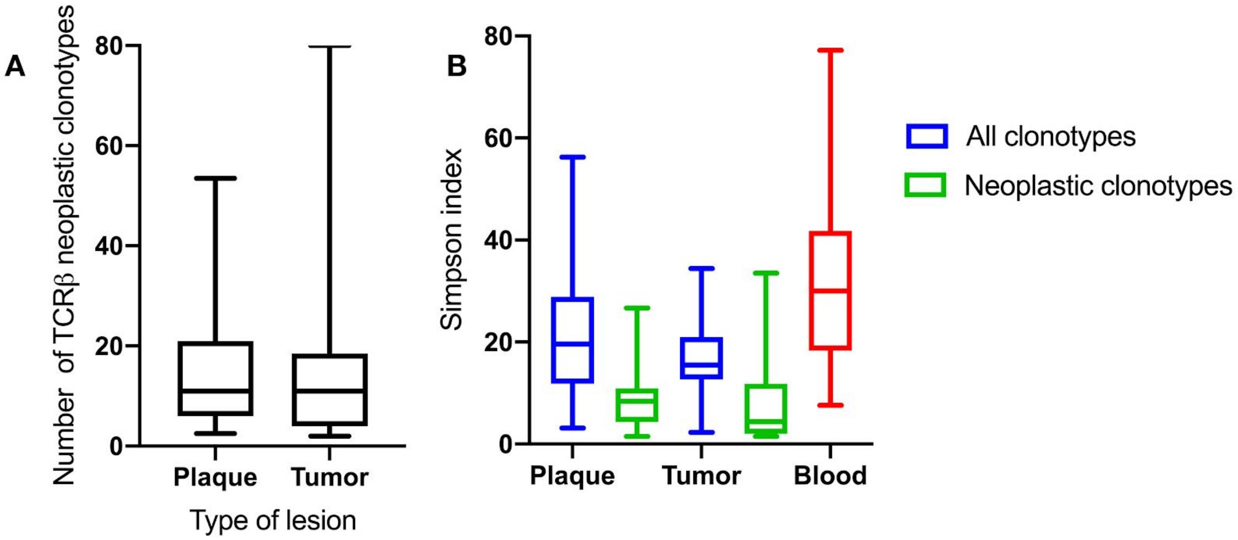


**Supplementary Figure S1: Comparison of the number of neoplastic TCRβ clonotypes and the Simpson diversity in different stages of skin lesions and blood.** Box plots represent median value with 25th-75th percentiles; the whiskers show the largest value within 1.5 times the interquartile range below the 25th percentile or above the 75th percentile. (A) represents the number of neoplastic TCRβ clonotypes in stages of skin lesions (plaque and tumor). (B) Indicates the simpson diversity index of neoplastic TCRβ clonotypes and all TCRβ clonotypes in skin lesions and blood.

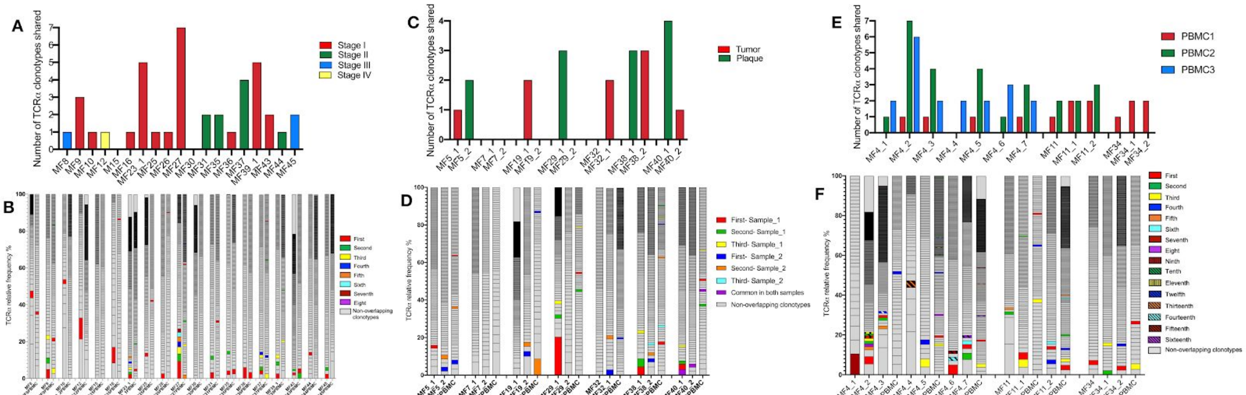


**Supplementary Figure S2: Shared neoplastic TCRα clonotypes in the skin and the peripheral blood.** The sequences of the TCRα clonotypes in the blood were matched with the sequences of the top 10 neoplastic TCRα clonotypes identified in the corresponding skin sample to identify the neoplastic clonotypes in the circulation. The number and frequency of those shared neoplastic clonotypes are shown separately for the three groups of patients as defined in Fig 1: A, B: 19 patients with a single biopsy, C-F: patients with multiple skin biopsies, of whom in 7 patients the biopsies were obtained at a single time point (C, D) whereas 3 patients were sampled longitudinally (E, F). In B, D, F the first ranking shared clonotype in the skin is indicated in red and the subsequent shared clonotypes are color-coded as indicated in the legend. The non-overlapping clonotypes are indicated in gray.

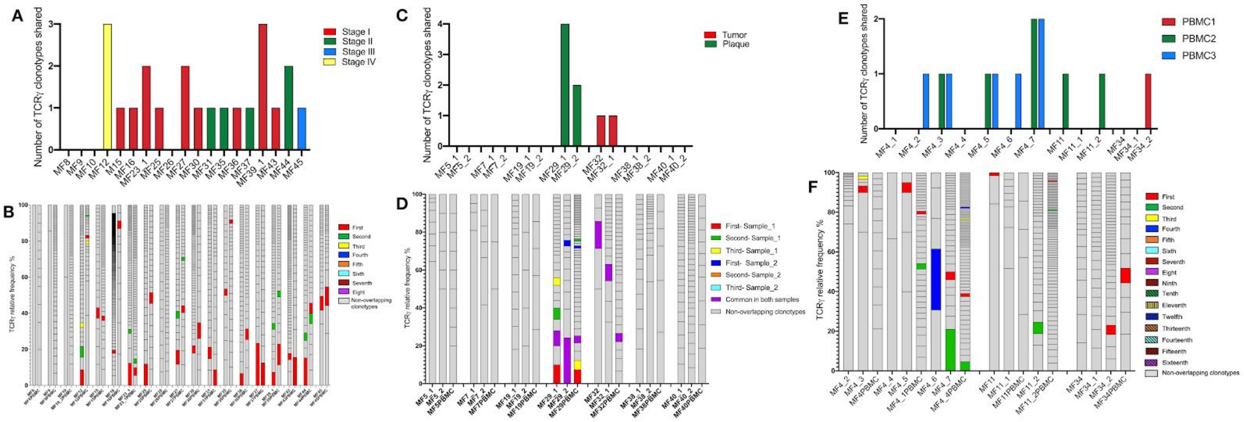


**Supplementary Figure S3: Shared neoplastic TCRɣ clonotypes in the skin and the peripheral blood.** The sequences of the TCRɣ clonotypes in the blood were matched with the sequences of the top 10 neoplastic TCRɣ clonotypes identified in the corresponding skin sample to identify the neoplastic clonotypes in the circulation. The number and frequency of those shared neoplastic clonotypes are shown separately for the three groups of patients as defined in Fig 1: A, B: 19 patients with a single biopsy, C-F: patients with multiple skin biopsies, of whom in 7 patients the biopsies were obtained at a single time point (C, D) whereas 3 patients were sampled longitudinally (E, F). In B, D, F the first ranking shared clonotype in the skin is indicated in red and the subsequent shared clonotypes are color-coded as indicated in the legend. The non-overlapping clonotypes are indicated in gray.

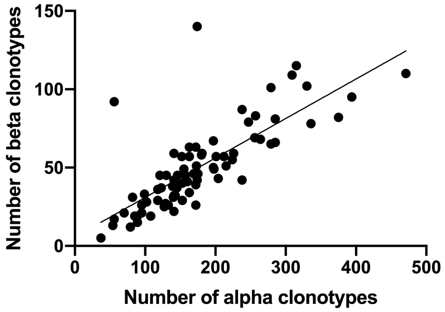


**Supplementary Figure S4: Correlation between the number of TCRα and TCRβ clonotypes in the skin lesions of MF.** Pooled data from all 46 samples. The regression coefficient is 0.786. Note that all clonotypes are shown.

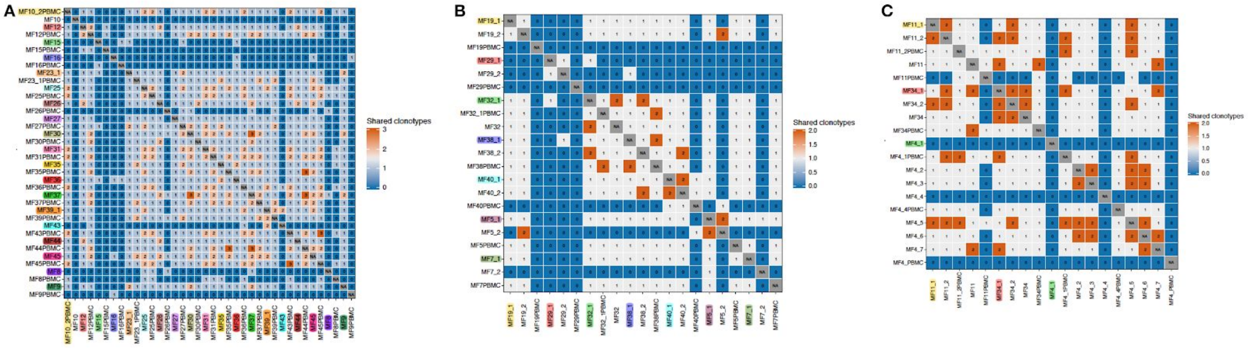


**Supplementary Figure S5: Neoplastic TCRβ clonotypes shared in the skin and the peripheral blood.** Top 10 most frequent TCRβ clonotypes from skin and peripheral blood were compared for shared CDR3aa sequences. The number of those shared neoplastic clonotypes are shown separately for the three groups of patients as defined in Fig 1: (A) 19 patients with a single biopsy, (B) patients with multiple skin biopsies, of whom in 7 patients the biopsies were obtained at a single time point whereas 3 patients were sampled longitudinally (C).

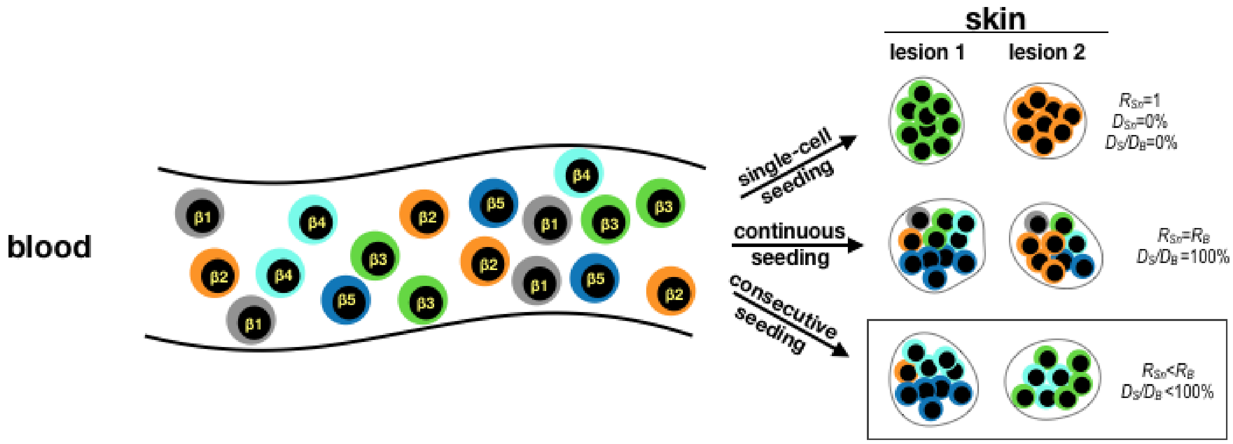


**Supplementary Figure S6: Models of cancer seeding.** The simplest scenario is the **single-cell seeding (upper row)** when single neoplastic clonotypes (as detected by unique TCRβ sequences β1-β1 marked with different colors enter the target tissue (skin) from the circulation and develop into the lesions. In this case, each lesion represents a single clone (clonotypic richness of any single *n*-lesion $R_{Sn}$=1) and the Simpson diversity index (the probability of detecting two different clonotypes in a single skin lesion, $D_{Sn}$) is 0%. Note, that discrete lesions may originate from different clones. During **continuous seeding (middle row)**, all malignant clonotypes will be detectable in any given skin lesion, either because the metastasis is mediated by clusters of malignant cells or because cooperation between all malignant clones is required for tumor growth. In this situation clonotypic richness of each lesion will be the same and equal to the clonotypic richness of all lesions combined ($\sum R_{Sn}$) and average richness of the lesions ($R_S$) and to the neoplastic cells in the blood

($R_{Sn}=R_{Sn+1}\ldots=\sum R_{Sn}=R_S=R_B$). The **consecutive seeding** is an intermediate situation when only a portion of circulating clones seeds the lesions (lower row, frame). Here, the clonotypic richness of any single skin lesion will be lower that the richness of all lesions combined or the richness of the entire repertoire of clonotypes in the blood ($R_{Sn}<\sum R_{Sn}\leq R_B$) and the average Simpson index of skin lesions ($D_S$) will be lower that the entire blood ($D_S<D_B$, $D_S/D_B<100\%$). This model is compatible with the data reported in this study. Note, that the richness and Simpson index of the neoplastic clonotypes in the blood ($R_B$, $D_B$) cannot be determined precisely experimentally because of insufficient sampling due to a low frequency of circulating, neoplastic clones. The model is modified from Ref[39] For simplicity, the traffic of cancer clones is marked as unidirectional movement from the blood to the skin, however it is likely that a recirculation of malignant clones between the blood and the tumors takes place as well.[36,52]

**Supplementary Table S1:** Patient characteristics and samples included in the study

| Patient ID ( age [years], sex [M-male, F-female]) | Sample ID | Lesion type | Diagnosis and stage[1] | Stage progression[2] | TTP[3] or PFS[4] (months)[3] |
|---|---|---|---|---|---|
| MF4 (69, M) | MF4_1P | Plaque | Mycosis Fungoides IIB | no | 30 |
| | MF4_2T | Tumor | | | |
| | MF4_3P | Plaque | | | |
| | MF4_4T | Tumor | | | |
| | MF4_5P | Plaque | | | |
| | MF4_6T | Tumor | | | |
| | MF4_7T | Tumor | | | |
| MF5 (44, F) | MF5_1T | Tumor | Folliculotropic Mycosis Fungoides IIB | no | 31 |
| | MF5_2P | Plaque | | | |
| MF7 (62, M) | MF7_1T | Tumor | Mycosis Fungoides IVA2 | no | 17 DOD[5] |
| | MF7_2P | Plaque | | | |
| MF8 (54, F) | MF8P | Plaque | Folliculotropic Mycosis Fungoides IIIB | yes | 3 |
| MF9 (42, F) | MF9P | Plaque | Mycosis Fungoides IA | no | 29 |
| MF10 (56, M) | MF10P | Plaque | Mycosis Fungoides IB | no | 29 |
| MF11 (56, M) | MF11T | Tumor | Mycosis Fungoides IIB | no | 29 |
| | MF11_1P | Plaque | | | |
| | MF11_2P | Plaque | | | |
| MF12 (66, M) | MF12P | Plaque | Mycosis Fungoides IVA2 | no | 27 |

| | | | | | |
|---|---|---|---|---|---|
| MF15 (65, M) | MF15P | Plaque | Mycosis Fungoides IB | no | 27 |
| MF16 (68, M) | MF16P | Plaque | Mycosis Fungoides IB | no | 26 |
| MF19 (74, M) | MF19_1T | Tumor | Mycosis Fungoides IIB | no | 13<br>DOD[5] |
| | MF19_2P | Plaque | | | |
| MF23_1 (69, F) | MF23_1P | Plaque | Mycosis Fungoides, IA | no | 22 |
| MF25 (48, F) | MF25P | Plaque | Mycosis Fungoides IB | no | 21 |
| MF26 (76, M) | MF26P | Plaque | Mycosis Fungoides IB | yes | 6 |
| MF27 (71, M) | MF27P | Plaque | Mycosis Fungoides IA | no | 16 |
| MF29 | MF29_1P | Plaque | Mycosis Fungoides IA | no | 15 |
| | MF29_2P | Plaque | | | |
| MF30 (62, M) | MF30P | Plaque | Mycosis Fungoides IB | yes | 2 |
| MF31 (67, M) | MF31T | Tumor | Folliculotropic Mycosis Fungoides IIB | no | 19 |
| MF32 | MF32T | Tumor | Mycosis Fungoides IVA | no | 27 |
| | MF32_1T | Tumor | | | |
| MF34 | MF34T | Tumor | Mycosis Fungoides IIB | no | 20 |
| | MF34_1P | Plaque | | | |
| | MF34_2T | Tumor | | | |
| MF35 (54, M) | MF35P | Plaque | Folliculotropic Mycosis Fungoides IIB | no | 11 |
| MF36 (64, M) | MF36P | Plaque | Mycosis Fungoides IA | no | 12<br>DOD[5] |

| | | | | | |
|---|---|---|---|---|---|
| MF37 (63, M) | MF37 | Tumor | Mycosis Fungoides IIB | no | 16 |
| MF38 | MF38_1P | Plaque | Mycosis Fungoides IIB | no | 15 |
| | MF38_2T | Tumor | | | |
| MF39 (71, M) | MF39_1P | Plaque | Mycosis Fungoides IB | yes | 12 DOD[5] |
| MF40 | MF40_1P | Plaque | Mycosis Fungoides IIB | no | 29 |
| | MF40_2T | Tumor | | | |
| MF43 (60, M) | MF43P | Plaque | Folliculotropic Mycosis Fungoides IA | no | 15 |
| MF44 (85, M) | MF44T | Tumor | Mycosis Fungoides IIB | no | 23 DOD[5] |
| MF45 (77, M) | MF45P | Plaque | Mycosis Fungoides IIIA | yes | 9 DOD[5] |

[1]established at the time of the biopsy; [2]observed between the time of the biopsy and last follow-up; [3]TTP - only for patients who progressed in stage during follow-up (months); [4]PFS - progression free survival, for the patients who remained in the same stage during follow-up (months); [5]DOD - dead of disease

**Supplementary Table 2:** CDR3aa sequences and the predicted V and J combinations for the TCR clonotypes that are shared between at least two skin biopsies (including across different MF patients) or skin biopsies and peripheral blood.

| CDR3 sequence | Samples | V usage | J usage |
|---|---|---|---|
| CDNNNDMRF | MF4_2, MF4_3, MF4_1PBMC, MF4_6, MF4_7, MF4_4PBMC, MF9, MF9PBMC, MF11_1, MF11PBMC, MF11_2PBMC, MF23_1, MF23_1PB, MF29_1, MF29PBMC, MF34_1, MF34PBMC, MF38_1, MF38PBMC, MF40_1, MF40_2, MF40PBMC | TRAV16 | TRAJ43 |
| CAG_AGA | MF4_2, MF4_3, MF4PBMC, | TRAV35 | TRAJ4 |

| | | | |
|---|---|---|---|
| | MF4_1PBMC, MF4_7, MF4_4PBMC, MF11_1, MF11PBMC, MF11_2, MF11_2PBMC, MF27, MF27PBMC, MF32_1, MF32_1PBMC, MF34_1, MF34PBMC, MF39_1, MF39PBMC, MF40_1, MF40PBMC, MF43, MF43PBMC | | |
| CGCENSGGSNYKLTF | MF5_2, MF5PBMC, MF11_2, MF11PBMC, MF11_2PBMC, MF16, MF16PBMC, MF19_1, MF19PBMC, MF27, MF27PBMC, MF37, MF37PBMC, MF38_1, MF38PBMC, | TRAV3 | TRAJ53 |
| CYEASHSVNTGTASKLTF | MF4_2, MF4_3, MF4_1PBMC, MF9, MF9PBMC, MF23_1, MF23_1PBMC, MF38_1, MF38PBMC | TRAV31 | TRAJ44 |
| CMIVGSPELTF | MF4_2, MF4_1PBMC, MF26, MF26PBMC, MF36, MF36PBMC, MF43, MF43PBMC | TRAV35 | TRAJ10 |
| CAGPCVFP*RREICYL_LNDALMWAKVNPFSPL | MF4_2, MF4_5, MF4_1PBMC, MF4_4PBMC, MF8, MF8PBMC, MF31, MF31PBMC | TRAV25 | TRAJ48 |
| CAV*G_LTNFS | MF5_2, MF5PBMC, MF11, MF11PBMC, MF11_2PBMC, MF27, MF27PBMC, | TRAV36 | TRAJ21 |
| IGEWELVLPPTK_LTRFLPAEQLPF | MF23_1, MF23_1PBMC, MF25, MF25PBMC, MF45, MF45PBMC | TRAV13-2 | TRAJ46 |
| CALVTAFLEPFF | MF4_7, MF4_1PBMC, MF4_4PBMC, MF27, MF27PBMC | TRAV39 | TRAJ12 |
| CGCDLHHRGSWK_VQGDEGLDKLHF | MF4_2, MF4_4PBMC, MF5_1, MF5PBMC | TRAV8-4 | TRAJ34 |
| CALRCLKSILNYNDHSIGTTF | MF4_2, MF4_1PBMC, MF37, MF37PBMC, | TRAV41 | TRAJ3 |
| CAVRSNWLECVRDGWEGGRYF*DGGSPCLEESI | MF35, MF35PBMC, MF39_1, MF39PBMC | TRAV21 | TRAJ5 |

| | | | |
|---|---|---|---|
| HQLHPQCA | | | |
| CAVKNTGLTAPC_SSSRYL PSRDF | MF4_5, MF4PBMC, MF4_1PBMC, MF4_4PBMC | TRAV3 | TRAJ50 |
| WTPY*E_F*ETLF | MF4_1, MF 4_4PBMC | TRAV8-3 | TRAJ8 |
| CVASGC*PRFPPLHSVSPL STPGAGTAPALIPQSS*SL RQTF | MF4_2, MF4_1PBMC | TRAV12-1 | TRAJ48 |
| CRE_IIF | MF4_3, MF4_1PBMC | TRAV7 | TRAJ30 |
| CAVRDTVTMRPL*L_PKFK RQPQQREAVF | MF4_4, MF4_4PBMC | TRAV1-2 | TRAJ51 |
| CSPEGPWPSLEPRT*VHLT *S_GIRIPEFSQPGVQNSR VRLL | MF4_6, MF4_4PBMC | TRAV34 | TRAJ24 |
| CAARHPPSWRQIIGSPRTE PLGVP_LPRPGQPGRRAA GGLLHLQHQLW | MF4_6, MF4_4PBMC | TRAV13-1 | TRAJ33 |
| CAVEG_EHLPF | MF4_4, MF4_4PBMC | TRAV36 | TRAJ5 |
| CALS_RR*G | MF9, MF9PBMC | TRAV19 | TRAJ4 |
| FERNAILFIESLKACFTCL FLKV*_EGVQQGCDRSS KN*NSFIYGHLIF | MF10, MF10_2PBMC | TRAV5 | TRAJ14 |
| CVWRQGVPPESPGLVRIS P_GTQKRKGQSREWGSD HSQQ | MF11, MF11_2PBMC | TRAV29 | TRAJ57 |
| CGSNRYSS*LFH*CQ**SLK WRSC_LSIDKVLTVFLFQG *IGSESSLVF | MF11_2, MF11_2PBMC | TRAV34 | TRAJ47 |
| CVCECS*CVYICVCVCVYV _VFVTDLIPVLSAGSILTC | MF12, MF12PBMC | TRAV10 | TRAJ58 |
| WQGLLLTGEERTSWERT | MF19_1, MF19PBMC | TRAV27 | TRAJ18 |
| SDSFRQSLLGSRRGRSSL SLAKSVSTTNIAG*ASVW | MF23_1, MF23_1PBMC | TRAV36 | TRAJ55 |
| CAL_AF | MF23_1, MF23_1PBMC | TRAV15 | TRAJ27 |
| CALSGTVAGF | MF27, MF27PBMC | TRAV16 | TRAJ16 |

| | | | |
|---|---|---|---|
| CALRDRVG | MF27, MF27PBMC | TRAV18 | TRAJ36 |
| YAVSGLIFSASLGFLQS_RKLCAYSGAGSYQLTF | MF27, MF27PBMC | TRAV8-4 | TRAJ28 |
| SAVGPAQDPPCHRQWGPCVHP_TLRPSGTRPPMWTGVWTGSPL | MF29_1, MF29PBMC | TRAV25 | TRAJ30 |
| CLVGLLMVTAHTTSPWLRVLIWRACRGIPGPMRASG | MF29_1, MF29PBMC | TRAV4 | TRAJ30 |
| CAGPCVFP*RREICYL_LNDALMWAKVNPFSPL | MF31, MF31PBMC | TRAV25 | TRAJ44 |
| CALRDRVGGTAARAQ_TNPGERRWGLGVSRF | MF31, MF31PBMC | TRAV18 | TRAJ14 |
| CCVGI_TNFS | MF32_1, MF32_1PBMC | TRAV28 | TRAJ21 |
| FMVNVTGELARQHYALSSL*PLHFFGQVKR**DL*L*LDARHTAS*FLKFFF | MF34, MF34_2, MF34PBMC | TRAV3 | TRAJ36 |
| CAGQHSAPQT_LQPVLKLAA | MF34_2, MF34PBMC | TRAV35 | TRAJ8 |
| CVVSEMFLSLLISKIFQGV*PQDITH_GPYAGGSLNRSN*EK*KTESKILIF | MF37, MF37PBMC | TRAV8-2 | TRAJ37 |
| VFYVSSHLGVPLSYVPALFSDG_LLLREGVALTVIVYQNLLYLTF | MF37, MF37PBMC | TRAV8-2 | TRAJ2 |
| CVVSGVSSGLLVPRLGIGWFEQTLGSGLI | MF38_2, MF38PBMC | TRAV10 | TRAJ4 |
| SASCTMRSISSLLRRPLSLVMVILLALPVLLSVAVTFRMPLASMSKVTSIW | MF38_2, MF38PBMC | TRAV3 | TRAJ33 |
| CAESTHCFSGTCILYPNLHLGLKPHSISFTF | MF38_2, MF38PBMC | TRAV5 | TRAJ13 |
| CAATTSGTYKYIF | MF39_1, MF39PBMC | TRAV21 | TRAJ40 |
| CVGVFQHGKVEIIANDQGN_HHPQLRGLHRHRAAHWGC | MF40_1, MF40PBMC | TRAV12-1 | TRAJ54 |
| *PGGCLSLFIRDWPRLNLL_GHWGLRLQALASSRPRGI | MF40_1, MF40PBMC | TRAV1-2 | TRAJ9 |

| | | | |
|---|---|---|---|
| CLLGDFPSLGLFLMWW*IHGS*RP_CGFERP*AGAPG*T*IYLLVPPF | MF44, MF44PBMC | TRAV40 | TRAJ22 |
| CAGPCSWTRY_AVSSHHLTF | MF45, MF45PBMC | TRAV25 | TRAJ46 |
| | | **TCR beta clonotypes** | |
| CAASRGC_AKNIQYF | MF4_2, MF4_3, MF4_5, MF4_1PBMC, MF4_6, MF4_7, MF4_4PBMC, MF5_1, MF5PBMC, MF11, MF11_1, MF11_2, MF11_2PBMC, MF12, MF12PBMC, MF23_1, MF23_1PBMC, MF25, MF25PBMC, MF26, MF26PBMC, MF27, MF27PBMC, MF30, MF30PBMC, MF31, MF31PBMC, MF32, MF32_1, MF32PBMC, MF34, MF34_1, MF34_2, MF34PBMC, MF35, MF35PBMC, MF36, MF36PBMC, MF37, MF37PBMC, MF38_1, MF38_2, MF38PBMC, MF39_1, MF39PBMC, MF40_1, MF40_2, MF40PBMC, MF44, MF44PBMC, MF45, MF45PBMC | TRBV18 | TRBJ2-4 |
| CVS_GVL | MF25, MF25PBMC, MF31, MF31PBMC, MF40_1, MF40PBMC | TRBV5-3 | TRBJ2-4 |
| CASSEATALHG | MF30, MF30PBMC, MF34_2, MF34PBMC, MF37, MF37PBMC | TRBV6-1 | TRBJ2-2P |
| CAKQLNEDCSKTQPCDHTKGLECNF | MF4_5, MF4_1PBMC, MF4_4PBMC | TRBV23-1 | TRBJ2-4 |
| GPGTRLLVLGERGLLGRGRGR_WVWFLRGVPGLCSGANVLTF | MF4_6, MF4_7, MF4_1PBMC | TRBV13 | TRBJ2-6 |
| CGGGARKTGQSPRQPRPGPSF | MF10, MF10_2PBMC, MF11_2, MF11_2PBMC | TRBV22-1 | TRBJ2-4 |
| CLARRQR_TTVVVF | MF4_5, MF4_1PBMC, MF35, MF35PBMC | TRBV5-6 | TRBJ2-2 |
| CELECIWPGSRNFECRRGISETVDKTAWKKTERC*SQTG | MF4_2, MF4_4PBMC | TRBV30 | TRBJ2-7 |
| CGRCSALP**RCGGRIPCS | MF4_2, MF4_4PBMC | TRBV8-2 | TRBJ1-5 |

| | | | |
|---|---|---|---|
| PRSS_NIVVEARGLVPTHTA*LVPQHF | | | |
| CAT_AR | MF4_3, MF4_1PBMC | TRBV27 | TRBJ2-2P |
| CAQQPVQSELVQRCQQLQSRLSTLKI_KRRGEFLP*EAQSHTLSFPVETFAAA | MF4_4, MF4_4PBMC | TRBV5-4 | TRBJ2-2P |
| CASCPH_VSCRRP | MF4_6, MF4_1PBMC | TRBV4-2 | TRBJ2-4 |
| CAS*LWW_*KSV**A | MF4_6, MF4_4PBMC | TRBV6-2 | TRBJ1-3 |
| *AAAWPAAAG_LRRVGWLAG | MF9, MF9PBMC | TRBV5-5 | TRBJ2-2P |
| FLRLLKYIRKLSG_QLRVSSSLNKPS | MF11, MF11PBMC | TRBV22-1 | TRBJ1-4 |
| CARVPRAV_NTGELFF | MF11, MF11_2PBMC | TRBV5-1 | TRBJ2-2 |
| CASSYSNQPQHF | MF12, MF12PBMC | TRBV19 | TRBJ1-5 |
| CAI*S*HPAS_KSSHAYSLFF | MF23_1, MF23PBMC | TRBV10-3 | TRBJ2-2 |
| CGSSEEGT | MF30, MF30PBMC | TRBV8-2 | TRBJ1-6 |
| CASSF | MF37, MF37PBMC | TRBV12-1 | TRBJ2-2 |
| CTTMR*QSR*VRARAGGRAAC_GGDFRLSMRFPAPGRSTEHF | MF38_1, MF38PBMC | TRBV5-6 | TRBJ2-7 |
| CASSLD*REETDTQYF | MF39_1, MF39PBMC | TRBV5-6 | TRBJ2-3 |
| CASSEPGRNQPQHF | MF39_1, MF39PBMC | TRBV6-1 | TRBJ1-5 |
| CASSQDTALQSHCIPVHKPPGSARKLQGSV_APAPRAPVSIP*WPLMEFQSVVQPASAPS | MF40_2, MF40PBMC | TRBV3-1 | TRBJ2-2 |
| | | **TCR gamma clonotypes** | |
| CAAWDYH_GWFKIF | MF4_1, MF4_7, MF4_4PBMC, MF12, MF12PBMC, MF27, MF27PBMC, MF29_1, MF29PBMC, MF30, MF30PBMC, MF31, MF31PBMC, MF32, MF32_1, MF32_1PBMC, MF35, MF35PBMC, MF36, MF36PBMC, | TRGV10 | TRGJP1 |

| | | | |
|---|---|---|---|
| | MF37, MF37PBMC, MF39_1, MF39PBMC, MF43, MF43PBMC, MF44, MF44PBMC, MF45, MF45PBMC | | |
| CATAAGLL_WL*SLLP* | MF4_3, MF4_5, MF4_7, MF4_1PBMC, MF4_4PBMC | TRGV5P | TRGJ1 |
| STAVGLA_SQAVQDN | MF4_2, MF4_4PBMC, MF11_2, MF11_2PBMC, MF27, MF27PBMC | TRGVB | TRGJP1 |
| CATAAGLLVVVVF | MF4_6, MF4_4PBMC, MF11, MF11_2PBMC, MF23_1, MF23_1PBMC | TRGV5P | TRGJP1 |
| CATWENR_GWFKIF | MF12, MF12PBMC, MF29_1, MF29PBMC | TRGV8 | TRGJP1 |
| CAAWDYGTGWFKIF | MF12, MF12PBMC, MF29_1, MF29_2, MF29PBMC, MF44, MF44PBMC | TRGV10 | TRGJP1 |
| CAAWDYTK_TTGWFKIF | MF15, MF15PBMC | TRGV10 | TRGJP1 |
| CATWVLP_GWFKIF | MF16, MF16PBMC | TRGV5 | TRGJP1 |
| CATRT_YKKLF | MF23_1, MF23_1PBMC, MF29_1, MF29PBMC | TRGV8 | TRGJ1 |
| CATWD_TRELF | MF25, MF25PBMC | TRGV2 | TRGJ1 |
| CACWIRH_GDWIKTF | MF29_2, MF29PBMC | TRGV11 | TRGJP2 |
| STAVGLA_KSGSSR* | MF34_2, MF34PBMC | TRGVB | TRGJP1 |
| CATWDG_YYKKLF | MF39_1, MF39PBMC | TRGV4 | TRGJ1 |
| CAAWDYGTGWFKIF | MF44, MF44PBMC | TRGV10 | TRGJP1 |
